# Mass cytometry and artificial intelligence define CD169 as a specific marker of SARS-CoV2-induced acute respiratory distress syndrome

**DOI:** 10.1101/2020.09.22.307975

**Authors:** M. Roussel, J. Ferrant, F. Reizine, S. Le Gallou, J. Dulong, S. Carl, M. Lesouhaitier, M. Gregoire, N. Bescher, C. Verdy, M. Latour, I. Bézier, M. Cornic, S. Leonard, J. Feuillard, V.K. Tiwari, J.M. Tadié, M. Cogné, K. Tarte

**Author notes:** These authors contributed equally to this work (M. Roussel, J. Ferrant). These authors contributed equally to this work (M. Cogné, K. Tarte). Correspondence to (M.R.); (K.T.).

## Abstract

Acute respiratory distress syndrome (ARDS) is the main complication of COVID-19, requiring admission to Intensive Care Unit (ICU). Despite recent immune profiling of COVID-19 patients, to what extent COVID-19-associated ARDS specifically differs from other causes of ARDS remains unknown, To address this question, we built 3 cohorts of patients categorized in COVID-19^neg^ARDS^pos^, COVID-19^pos^ARDS^pos^, and COVID-19^pos^ARDS^neg^, and compared their immune landscape analyzed by high-dimensional mass cytometry on peripheral blood followed by artificial intelligence analysis. A cell signature associating S100A9/calprotectin-producing CD169^pos^ monocytes, plasmablasts, and Th1 cells was specifically found in COVID-19^pos^ARDS^pos^, unlike COVID-19^neg^ARDS^pos^ patients. Moreover, this signature was shared by COVID-19^pos^ARDS^neg^ patients, suggesting severe COVID-19 patients, whatever they experienced or not ARDS, displayed similar immune dysfunctions. We also showed an increase in CD14^pos^HLA-DR^low^ and CD14^low^CD16^pos^ monocytes correlated to the occurrence of adverse events during ICU stay. Our study demonstrates that COVID-19-associated ARDS display a specific immune profile, and might benefit from personalized therapy in addition to standard ARDS management.

**One Sentence Summary:** COVID-19-associated ARDS is biologically distinct from other causes of ARDS.

## Introduction

The SARS-Coronavirus-2 (SARS-CoV-2) virus has currently affected more than 30 million people worldwide, requiring admission to Intensive Care Unit (ICU) for more than 2 million patients (*1*). Whereas most patients exhibit mild-to-moderate symptoms, acute respiratory distress syndrome (ARDS) is the major complication of the coronavirus disease 2019 (COVID-19) (*2*, *3*), leading to prolonged ICU stays, and high frequency of secondary complications, notably cardiovascular events, thrombosis, pulmonary embolism, and stroke (*1*, *4*). The immune system plays a dual role in COVID-19, contributing to both virus elimination and ARDS development. Excessive inflammatory response has been proposed as the leading cause of COVID-19-related clinical complications, thus supporting intensive efforts to better understand the specificities and mechanisms of SARS-CoV-2-induced immune dysfunction (*5*, *6*). Moreover, even if antiviral strategies, such as those provided by remdesivir or convalescent plasma, can lower the viral burden, no antiviral treatment has yet been able to prevent the evolution of some patients towards deregulated inflammation and critical respiratory complications. Recent data however suggest a benefit of corticosteroids in lowering overall mortality in COVID-19 patients with moderate disease, severe disease, and ARDS (*7*). However, steroid therapy could be harmful in some specific ARDS etiologies, such as in influenza-associated ARDS (*8*). A better understanding of the etiology-specific immune dysfunctions underlying ARDS development and severity is thus a major unmet need to design specific therapeutic strategy.

A number of high-resolution studies have recently concentrated on the determination of circulating markers that can distinguish severe from mild forms of COVID-19, providing a tremendous amount of data describing phenotypic and functional alterations in T cell, B cell, and myeloid cell subsets (*9*–*20*). In particular, CD14^pos^HLA-DR^low^, CD14^pos^CD16^pos^, CD14^low^CD16^pos^, and immature monocytes were demonstrated as increased among peripheral blood mononuclear cells (PBMCs) from critically ill COVID-19 patients (*11*, *16*, *18*, *21*–*23*). Various alterations of lymphoid cells have also been described, including a T-cell lymphopenia, predictive of patient outcome, a broad T-cell activation including Th1, Th2, and Th17, an alteration of B-cell and T-cell repertoires, and a strong increase of plasmablasts, most prominent in ARDS COVID-19 patients (*12*, *20*, *24*–*26*). Importantly, whereas non-COVID-19 ARDS is associated with a large panel of immune alterations, COVID-19 ARDS immune profiling was performed using healthy donors as a control, thus precluding any conclusions on whether reported immune alterations could be related to COVID-19 and/or ARDS status. Answering this question has potential to decipher whether ARDS induced by SARS-CoV-2 is mechanistically different from other ARDS etiologies.

To fill this gap, we performed a high-throughput mass cytometry approach on PBMCs obtained from 3 complementary cohorts of 12 COVID-19^neg^ARDS^pos^, 13 COVID-19^pos^ARDS^pos^, and 17 COVID-19^pos^ARDS^neg^ patients. We report common myeloid cell alterations in all COVID-19 patients, which are absent from non-COVID-19 ARDS patients. This includes in particular a strong increase of an unusual population of activated monocytes showing upregulated expression of CD169, associated with major COVID-19-specific alterations of T and B-cell compartments.

## Results

### Study population

Analyses were performed on a cohort of 63 cryopreserved PBMC samples isolated from 42 patients included in ICU (n = 36) or infectious standard ward (n = 6). The demographic characteristics of patients included are provided in Table 1. All patients but one were classified as severe at admission, requiring oxygen at a flow rate higher than 2 liters/min. ARDS was defined in accordance with international guidelines (*27*). Patients were classified in 3 groups: COVID-19^neg^ARDS^pos^ (n = 12, ARDS stages: 1 mild, 4 moderate, 7 severe), COVID-19^pos^ARDS^pos^ (n = 13, ARDS stages: 8 moderate, 5 severe), and COVID-19^pos^ARDS^neg^ (n = 17, including 11 from ICU and 6 from infectious standard ward). In the COVID-19^pos^ARDS^neg^, no statistical differences were noticed for immune cell abundance or phenotype between ICU and standard ward patients. Within the COVID-19^neg^ARDS^pos^ group, ARDS etiologies were bacterial pneumonia (n = 9), antisynthetase syndrome (n = 1), and unknown (n = 2). No corticoid therapy was started at the time of sampling. For 21 patients, a second blood sample obtained on day 7 after enrollment was studied (n = 7 for COVID-19^neg^ARDS^pos^, n = 8 for COVID-19^pos^ARDS^pos^, and n = 6 for COVID-19^pos^ARDS^neg^).

**Table 1:** Patients characteristics.

|  | COVID-19<sup>neg</sup> | COVID-19<sup>pos</sup> | COVID-19<sup>pos</sup> |
| --- | --- | --- | --- |
|  | ARDS<sup>pos</sup> | ARDS<sup>pos</sup> | ARDS<sup>neg</sup> |
| Patients D0/D7, n | 12/7 | 13/8 | 17/6 |
| Age, median (IQR) | 62 (48.2-66.7) | 59 (53.5-67.5) | 55 (46-67) |
| Male, n (%) | 7 (58) | 10 (77) | 12 (71) |
| ICU/Clinical ward, n | 12/0 | 13/0 | 11/6* |
| SAPS II, median (IQR) | 44.5 (29.2-59.2) | 33 (19.5-39.5) | 22 (13-28)* |
| Length of stay in ICU, median (IQR) | 11.5 (4.5-18.7) | 15 (11-54) | 2 (1-2)** |
| Length of stay in Hospital, median (IQR) | 18 (7-30.5) | 22 (15-62.5) | 9 (7.5-13) |
| **Comorbidities** |  |  |  |
| BMI, median (IQR) | 26.4 (19.5-28.4) | 28.6 (25-32) | 28.1 (22.3-32.1) |
| Chronic cardiovascular disease, n (%) | 1 (8.3) | 3 (23) | 1 (5.8) |
| Diabetes, n (%) | 2 (16.7) | 3 (23) | 1 (5.8) |
| Chronic respiratory disease, n (%) | 1 (8.3) | 0 (0) | 0 (0) |
| Chronic kidney disease, n (%) | 0 (0) | 2 (15.4) | 0 (0) |
| Cancer, n (%) | 3 (25) | 0 (0) | 0 (0) |
| **Severity criteria** |  |  |  |
| Maximal O<sub>2</sub> (L/min), median (IQR) | 10 (7.5-15) | 14 (9.2-15) | 3 (2-5) |
| Invasive ventilation, n (%) | 12 (100) | 13 (100) | 0 (0) |
| PaO<sub>2</sub>/FiO<sub>2</sub>, median (IQR) | 116.5 (75.2-161.9) | 106 (95.5-240) | 313 (218.5-340.3) |
| **Events occurring during follow up** |  |  |  |
| Thromboembolic, n (%) | 4 (33.3) | 4 (30.8) | 1 (5.8) |
| ICU-acquired infections, n (%) | 2 (16.7) | 7 (53.8) | 0 (0) |
| Septic shock, n (%) | 3 (25) | 2 (15.4) | 0 (0) |
| Renal failure, n (%) | 5 (41.7) | 8 (61.5) | 0 (0) |
| Deaths, n (%) | 4 (33.3) | 1 (7.7) | 0 (0) |

\*: all patients except 1 required O<sub>2</sub> at > 2 L/mn at admission; \*\*: For patients in ICU; n: number; IQR: interquartile range; SAPS II: simplified acute physiology score

### SARS-CoV2 induces phenotypic changes in circulating immune cells

To decipher the impact of SARS-CoV2 on circulating immune cells, we characterized PBMCs from COVID-19^pos^ *versus* COVID-19^neg^ patients at admission (D0) using two separate mass cytometry panels exploring myeloid and lymphoid subsets, respectively (Table S1). The full pipeline of analysis is depicted in fig. S1. First, we performed an unbiased discovery approach with CellCnn, a neural network-based artificial intelligence algorithm allowing analysis of single-cell data and detection of cells associated with clinical status (*28*–*30*). During training, CellCnn learns combinations of weights for each marker in a given panel that best discriminate between groups of patients. These weight combinations, called filters, can be used to highlight the specific profiles of cells associated with patient status. We identified the best-performing CellCnn filters for both the myeloid and the lymphoid panels highlighting a population of cells significantly enriched in COVID-19^pos^ patients as compared to COVID-19^neg^ patients (P < 0.0001 for both panels) (Fig. 1A). Projecting these cells on tSNE maps generated with either the myeloid or the lymphoid panels revealed that they fell into several distinct areas (Fig. 1B). The cells selected by the CellCnn filter on the myeloid panel showed high expression for CD169, CD64, S100A9, CD11b, CD33, CD14, and CD36 compared to background, while the cells selected by the CellCnn filter on the lymphoid panel showed high expression for CD38 and CXCR3 (Fig.1B and Fig. S2). This broad and unbiased approach showed that immune markers, in particular related to monocytes, segregated COVID-19^neg^ and COVID-19^pos^ patients.

**Fig. 1 :**
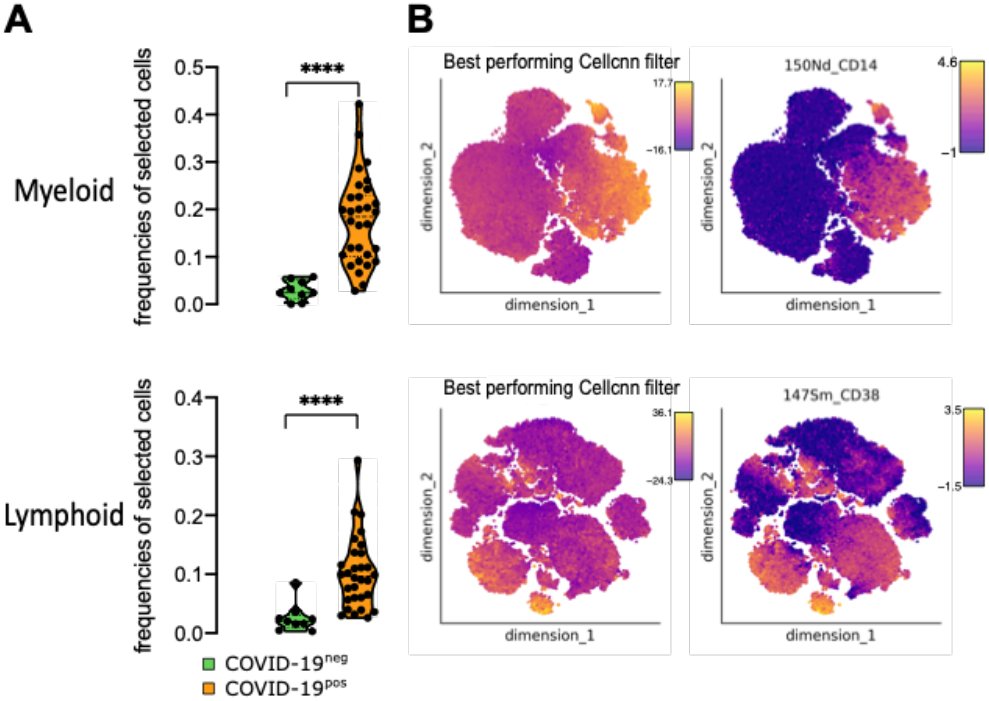
SARS-CoV2 induces specific phenotype of circulating immune cells. CellCnn analysis performed on single cells from myeloid (top) and lymphoid (bottom) panels on 39 samples at admission (Day 0) (COVID-19^neg^ [n = 9] and COVID-19^pos^ [n = 30]). **(A)** Frequencies of cells discovered by the best-performing CellCnn filter in COVID-19^neg^ (blue) and COVID-19^pos^ (orange) patients for each panel. Mann-Whitney tests, ****P < 0.0001. **(B)** Cells defined by the best-performing CellCnn filters enrichment shown on tSNE and representative markers for each panel (CD14 and CD38 [see additional markers in Fig. S2]).

### SARS-CoV2 induces CD169-expressing monocyte subsets

To investigate circulating monocyte heterogeneity and define consistent phenotypes, we used the FlowSOM algorithm. This approach led to the identification of 15 monocyte metaclusters from the myeloid panel (Fig. 2A). In particular, Mo30, Mo11, and Mo28 metaclusters were defined by higher expression of CD16 and lower expression of CD14, CD36, and CD64, corresponding to a non-classical monocyte phenotype. Mo21 and Mo22 were defined by the high expression of S100A9 and the low expression of CD36. Finally, Mo243 and Mo180 strongly expressed S100A9, CD169, and CD36. To assess the phenotypic changes in monocytes during SARS-CoV2 infection, we determined the frequencies of these metaclusters in each patient at admission and performed hierarchical clustering on these values (Fig. 2B). The upper branch of the hierarchical clustering included 20 COVID^pos^ (10 ARDS^neg^ and 10 ARDS^pos^) and 1 COVID^neg^ARDS^pos^ patient whereas the lower branch included 10 COVID^pos^ (7 ARDS^neg^ and 3 ARDS^pos^) and 11 COVID^neg^ARDS^pos^ (chi-square = 0.001) (Fig. 2B). We then analyzed the abundance of individual metaclusters and identified only 4 metaclusters out of 15 as differentially represented between the 3 groups of patients (Fig. 2C and Fig. S3). In particular, within ARDS^pos^ patients, Mo11 and M181 were less abundant in COVID-19^pos^ patients (P < 0.01 and P < 0.05, respectively), while Mo243 and Mo180 were more abundant (P < 0.05 and P < 0.001) (Fig. 2C). No differences were detected within COVID-19^pos^ groups (ARDS^pos^ versus ARDS^neg^) (Fig. 2C). Interestingly, Mo243 and Mo180 were both enriched in cells highly expressing CD169, CD64, CD36, and CD14 (Fig. 2A and 2D). Additionally, Mo22 was present only in some COVID^pos^ patients and also expressed CD169 (Fig. 2B). Taken as a whole, Mo243, Mo180, and Mo22 clusters were highly enriched in COVID-19^pos^ patients when compared to COVID-19^neg^ patients (P < 0.0001), with no difference regarding the ARDS status (Fig. 2E). Accordingly, CD169 was differentially expressed in COVID-19^pos^ versus COVID-19^neg^ patients (P < 0.001) (Fig. 2E). As a whole, our study including COVID-19 and non-COVID-19 critically ill patients suggest a specificity of CD169 expression in COVID-19 patients, and greatly extend previous scRNAseq data showing an expansion of CD169-expressing monocytes in COVID-19 patients compared to healthy donors (Fig. 2F) (*20*).

**Fig. 2 :**
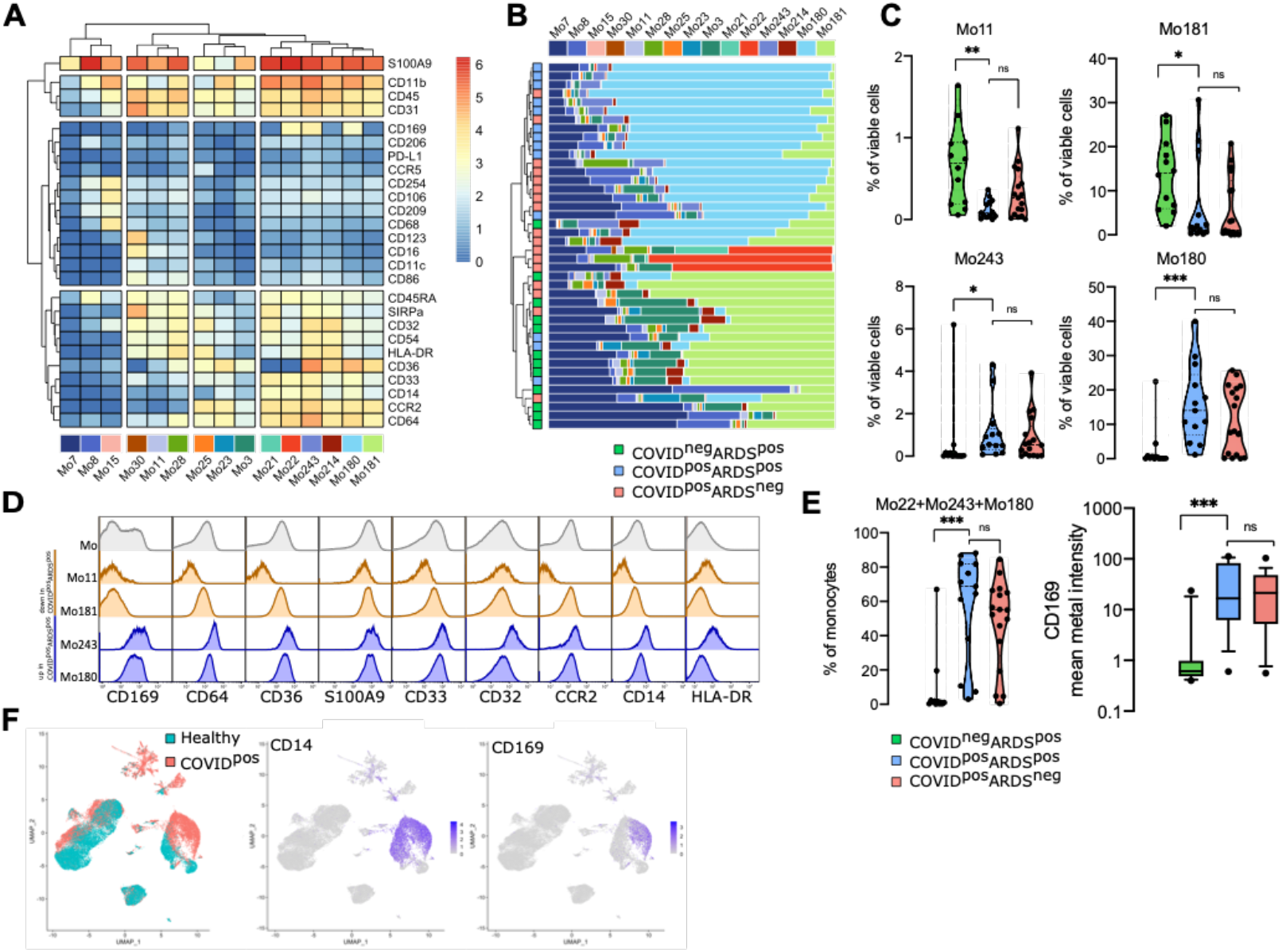
CD169 monocytes are enriched in SARS-CoV2 infected patients. **(A)** Heatmap of the 15 monocyte metaclusters defined after FlowSOM analysis. **(B)** Relative abundance of metaclusters among monocytes for each patient and hierarchical clustering of COVID-19^neg^ARDS^pos^ (n=12, green), COVID-19^pos^ARDS^pos^ (n=13, blue), and COVID-19^pos^ARDS^neg^ (n=17, red). **(C)** Abundance of metaclusters differentially expressed between groups, among singlet cell analyzed. **(D)** Expression of the corresponding markers (mean metal intensity) for background (gray), Mo11 and Mo181 (orange), and Mo243 and Mo180 (blue) metaclusters. **(E)** Abundance of Mo22, Mo180, and Mo243 and expression of CD169 (Bow and Whiskers with 10 and 90 percentile). **(F)** UMAP from scRNAseq of COVID-19 patients (COVID-19) and healthy donors (healthy) highlighting CD14 and CD169 expression (*20*). Kruskal-Wallis test with Dunn’s multiple comparison correction, *P < 0.05, **P < 0.01, ***P < 0.001.

### Monocyte metacluster enrichment in COVID-19 is correlated with a specific increase of effector memory T cells and plasma cells

To define a more global immune pattern and the relationship between immune cells in the context of the SARS-CoV2 infection, we sought for correlation between frequencies of clusters of T, NK, B, and plasma cells (n = 136 clusters from the lymphoid panel, fig. S4) and the 4 monocyte metaclusters (Mo11, Mo181, Mo243, and Mo180) previously described. This analysis identified 70 clusters with significantly correlated variations (P < 0.05) (Fig. S5). To strengthen the relevance of these correlations, we restrained further analysis to the 29 strongest relationships (R > 0.5 or < −0.5 and P < 0.01) between Mo180 or Mo243 (the two metaclusters enriched in COVID-19 patients) and other immune cell subsets (Fig. 3A and Table S2). As expected, Mo180 and Mo243 clusters were correlated (R = 0.93). Moreover, they were positively correlated with 18 clusters of T (n = 6), NK (n = 10), and plasma cells (n = 2), and inversely correlated with 11 clusters of T (n = 9), and NK cells (n = 2) (Fig. 3A). Among positively correlated clusters, plasmo_183 and plasmo_198 similarly expressed CD38, CD44, and CD27, whereas plasmo_183 was high for Ki-67 and HLA-DR, corresponding to an early plasma cell phenotype (Fig. 3B). NK cells were all marked by CD7 and T-bet expression, NK_209 being CD8^high^, and NK_241 and NK_197 displaying a Ki-67^high^ proliferating phenotype. The related T8_147 and T8_161 clusters exhibited a CD45RA^high^CD45RO^low^CCD7^low^CD27^low^Tbet^high^CD38^high^ effector phenotype. Few T4 clusters were positively correlated with Mo180 and Mo243, among them T4_106 displayed an effector memory proliferating phenotype (Ki-67^high^CD45RA^low^CCR7^low^CD45RO^high^CD27^high^ and CTLA4^high^PD1^high^). T4_25 was also marked by an effector memory phenotype (CD45RA^low^CCR7^low^CD45RO^pos^) and displayed a CD27^low^CD127^pos^CCR6^pos^CxCR3^neg^CD161^pos^ Th17 profile (Fig. 3B). Conversely, some T4 clusters were inversely correlated with Mo_180 and Mo_243, in particular clusters T4_6, T4_20, and T4_34, all three corresponding to naïve cells (CD45RA^high^CD45RO^low^CCR7^high^), and T4_59 expressing a Th2 phenotype (CCR4^high^). We then compared the abundance of these 29 lymphoid clusters correlated with Mo180 and Mo243 and highlighted the 22 differentially represented lymphoid clusters between the three groups of patients (P < 0.05) (Fig. 3C and Fig. S6). Only 7 clusters of CD4 T cells, and 2 clusters of CD8 T cells were at lower abundance in COVID-19^pos^ARDS^pos^ patients compared to COVID-19^neg^ARDS^pos^ patients. As previously discussed, T4_6, T4_20, and T4_34 corresponded to naïve cells, whereas within the effector memory cells, T4_7 and T4_45 were CD127^low^, T4_24, T8_99, and T8_113 were CD127^high^, and T4_59 was CCR4^high^. Conversely, 13 clusters were enriched in COVID-19^pos^ARDS^pos^ compared to COVID-19^neg^ARDS^pos^ including: i) CTLA4^high^PD1^high^ effector memory activated CD4 Tcells (T4_106); ii) Tbet^high^ Th1-like CD8 effector phenotype (T8_146, T8_147, and T8_161); iii) cytotoxic mature CD16^pos^CD56^low^CD7^pos^Tbet^pos^CD127^neg^ NK cells (NK_209, NK_241, NK_242, and NK_244) with in particular proliferating Ki-67^high^ NK cells (NK_241); and iv) proliferating plasmablasts (plasmo_183) and mature plasma cells (plasmo_198) (Fig. 3B and Fig. 3C). Of note, no cluster was differentially expressed between COVID-19^pos^ARDS^pos^ and COVID-19^pos^ARDS^neg^ groups (Fig. 3C and Fig. S6). Then, to explore the whole immune profile and define relationship with groups of patients, we performed correspondence analysis (CA) using, as a variable, the abundance of the myeloid (n = 4) and the lymphoid (n = 22) clusters differentially expressed between groups of patients (Fig. 3D). CA was developed to analyze frequency tables and visualize similarities between patients and co-occurrence of cell subsets (*31*). The first and second dimension of the correspondence analysis explained 80.5 % and 13.5 % of the difference, respectively (Fig. 3D). The top-ten cell populations explaining the difference between COVID^pos^ and COVID^neg^ patients were Mo243, Mo180, T8_146, NK_244, and T8_161 being increased and Mo181, T4_6, Mo11, T8_99, and T4_45 being decreased in COVID^pos^. Altogether, these subsets corresponded to an increase in inflammatory monocytes (CD169^high^ CD64^high^), Tbet^high^ Th1-like CD8 T cells, and mature NK cells and a decrease in naïve T4 cells and effector memory T4 and T8 cells. Interestingly, only the first dimension of the correspondence analysis segregated COVID-19^pos^ARDS^pos^ from COVID-19^neg^ARDS^pos^ (P < 0.001) and no statistical differences were found between COVID-19^pos^ARDS^pos^ and COVID-19^pos^ARDS^neg^ (Fig. 3D).

**Fig. 3 :**
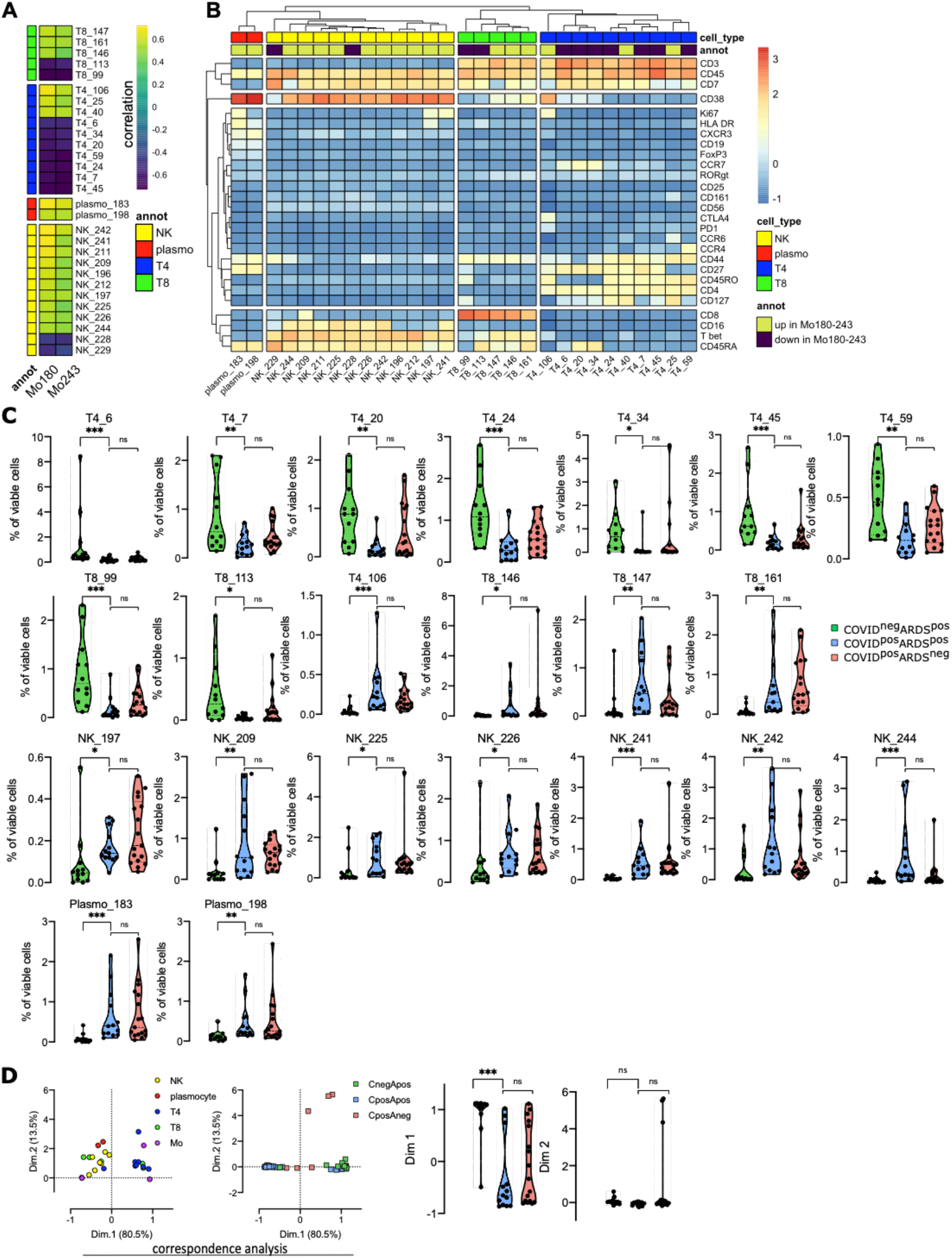
Monocyte clusters enriched in COVID-19 are correlated with effector memory T cells and plasma cells. **(A)** Correlation between Mo180 and Mo243 and lymphoid clusters (see heatmap for all lymphoid clusters and markers in Fig. S4) from all patients at D0 (COVID-19^neg^ARDS^pos^ [n=12], COVID-19^pos^ARDS^pos^ [n=13], and COVID-19^pos^ARDS^neg^ [n=17]. Only strong correlations (Spearman R > 0.5 or R < −0.5 and P < 0.01) are shown (see all significant correlations [P < 0.05] in fig. S5 and table S2). **(B)** Heatmap showing marker expression for the lymphoid clusters (Spearman R > 0.5 or R < −0.5 and P < 0.001) strongly correlated with Mo180 and Mo243 (see heatmap for all clusters and markers in Fig. S4). **(C)** Abundance of lymphoid clusters differentially expressed between groups, among singlet cells analyzed. Kruskal-Wallis test with Dunn’s multiple comparison correction, *P < 0.05, **P < 0.01, ***P < 0.001 [see all clusters in Fig. S6]). **(D)** Two first dimensions of correspondence analysis accounting for 84 % of the association between immune clusters differentially expressed between groups (n= 4 monocyte- and n=22 lymphoid-clusters), and patients. For clarity, patients and immune cells are shown on 2 different plots. Dimensions 1 and 2 coordinates are compared between groups of patients. Kruskal-Wallis test with Dunn’s multiple comparison correction, ****P < 0.0001.

### Evolution of immune cell clusters between D0 and D7 in COVID-19 patients defines high-risk clinical grade

We next performed mass cytometry analysis for 21 patients at day 7 of hospitalization, including 7 COVID-19^neg^ARDS^pos^, 8 COVID-19^pos^ARDS^pos^, and 6 COVID^pos^ARDS^neg^ patients, in order to follow up the kinetic of PBMC phenotypic alterations. The 42 samples (21 at day 0 and 21 at day 7) were parsed by correspondence analysis using, as a variable, the abundance of myeloid and lymphoid clusters (Fig. 4A). The first and second dimensions of the correspondence analysis explained 85.1 % and 9 % of the differences acquired between D0 and D7. The first dimension captured the difference between D0 and D7 only for COVID-19^pos^ARDS^pos^ (P < 0.01) (Fig. 4A). Because of the limited number of samples, only a trend was observed for COVID^pos^ARDS^neg^ (P = 0.062). The top-five enriched populations explaining the differences between D0 and D7 for COVID-19^pos^ARDS^pos^ patients were Mo11, Mo181, T8_113, T4_34, and NK_197, corresponding to an enrichment in non-classical monocytes (CD14^low^CD16^high^CD64^low^CD36^low^S100A9^high^), in M-MDSC-like (HLA-DR^low^S100A9^high^), in effector memory CD127^high^ T8 cells, in T4 naïve cells, and in Ki-67^high^ proliferating NK cells. These 5 cell subsets were integrated in an immune score combining their fold change between D0 and D7. To define the relevance of this immune score in discriminating COVID-19 patients with unfavorable prognosis, we built a clinical score as the sum of events occurring during ICU stay (thromboembolic, ICU-acquired infection, septic shock, renal failure, and deaths) (Table 1). Interestingly, both the clinical and the immune scores were found correlated in severe COVID-19 patients, irrespectively of their ARDS status (Spearman R = 0.58; P < 0.05) (Fig. 4B).

**Fig. 4 :**
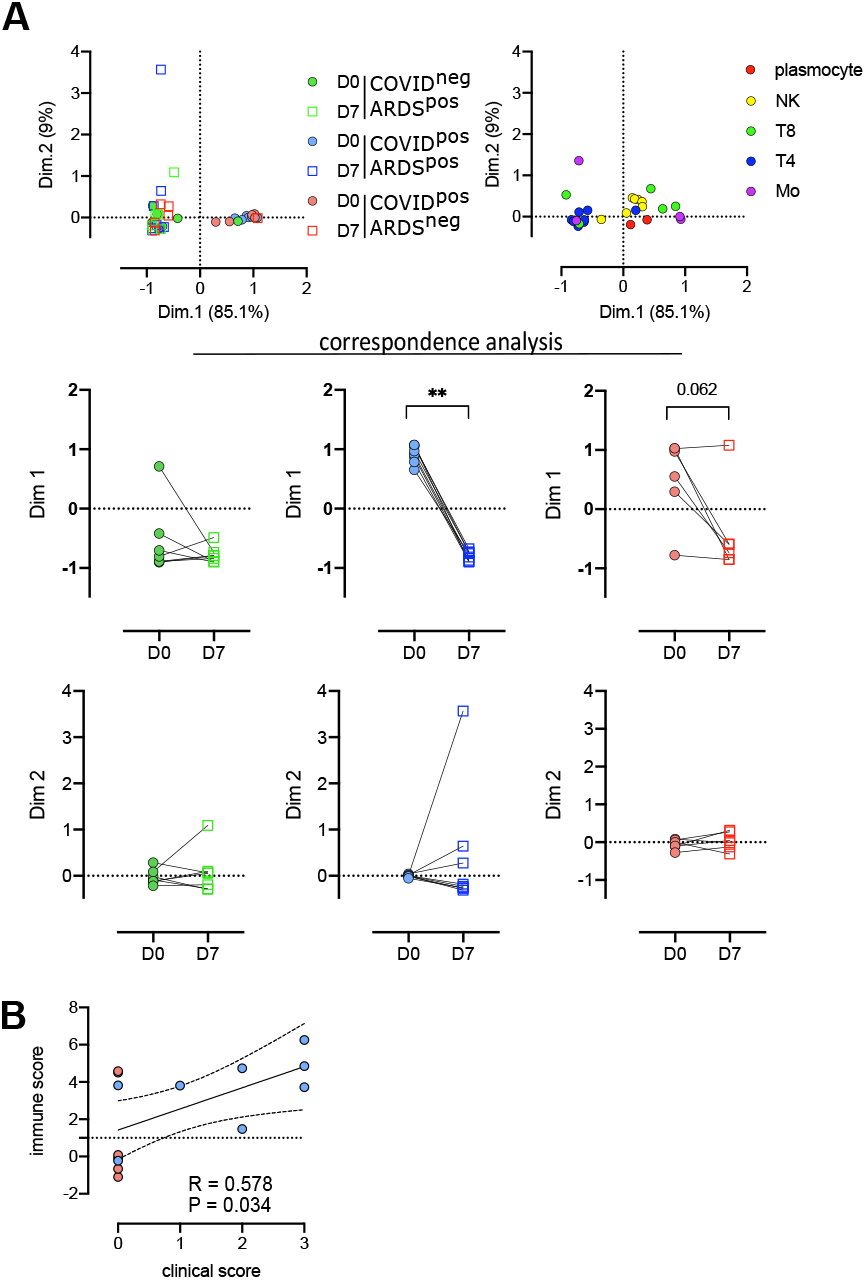
Evolution of immune cell subsets between D0 and D7, defines high-risk clinical grade COVID-19 patients. **(A)** Two first dimension of correspondence analysis accounting for 94.1% of the association between immune clusters differentially expressed between groups (n= 4 monocyte and n=22 lymphoid clusters), and patients for which a follow-up of 7 days was available (COVID-19^neg^ARDS^pos^ [n=7], COVID-19^pos^ARDS^pos^ [n=8], and COVID-19^pos^ARDS^neg^ [n=6]). For clarity, patients and immune cells are shown on 2 different plots. Dimensions 1 and 2 coordinates were compared between D0 and D7 for each group of patients. Wilcoxon matched-pairs signed rank tests, **P < 0.01. **(B)** Spearman correlation between immune and clinical score for COVID-19^pos^ patients (ARDS^pos^ [n=8] and ARDS^neg^ [n=6]).

## Discussion

Immune response to COVID-19 infection has been recently intensively studied at transcriptomic and proteomic levels. However, most studies focused on either the lymphoid (*15*, *17*, *19*) or the myeloid compartments (*9*, *16*, *18*), and only few performed a wide analysis of the circulating immune landscape (*10*, *12*, *13*, *20*, *32*), thus precluding the definition of complex patterns of immune parameter alterations associated with COVID-19 severity or physiopathology. Moreover, these works were designed to identify differences in immune cell subset frequencies between COVID-19 patients and healthy donors, and eventually correlated with the severity of the disease, but did not include severe non-COVID-19 patients as controls, although critically ill patients were largely demonstrated previously to display immune reprogramming (*33*). ARDS is a major adverse event occurring during ICU stay, leading to an overall mortality rate of 40 % to 60 %. Whether COVID-19 associated ARDS is clinically and biologically similar to other causes of ARDS remains controversial (*34*, *35*). To address this point, we characterized for the first time, by mass cytometry, the immune landscape in COVID-19-associated ARDS compared to other causes of ARDS. We demonstrated that an increase of CD169^pos^ monocytes, correlated with specific changes of T, plasma, and NK cell subsets, defines COVID-19-associated ARDS and is not found for bacteria-associated ARDS, suggesting a COVID-19 specific immune reprogramming.

The amplification of CD169^pos^ circulating monocytes has already been highlighted in the context of COVID-19 (*11*, *18*, *36*, *37*), and is reminiscent of other inflammatory conditions found in viral infections, such as with Human Immunodeficiency Virus or Epstein-Barr Virus, in which the CD169 sialoadhesin is induced in an IFN-dependent manner on the surface of circulating monocytes (*38*, *39*). Consistent with the inflammatory response, we showed that the accumulation of CD169^pos^ monocytes in COVID-19^pos^ patients is positively correlated with an increase of plasmablasts and mature plasma cells, Th1-like CD8 effector T cells, cytotoxic mature NK cells, and activated CD4 effector memory T cells displaying a CTLA4^high^PD1^high^ phenotype. CD169^pos^ activated monocytes were detected in mild disease (*18*), and were proposed to rise rapidly and transiently in patients with COVID-19, in association with a high expression of IFNγ and CCL8 (*11*). This could be due to the transient nature of this monocytic population, either losing CD169, being short-lived, or being recruited into tissues as CD169^pos^ macrophages, as suggested by the high expression of CCR2 on Mo243 and Mo180, the two monocyte subsets identified here in COVID-19 patients, and the local inflammation and lung tissue destruction mediated by monocyte-derived macrophages in severe cases of SARS-CoV2 infections (*40*, *41*). Interestingly, we also found an upregulation of cytoplasmic S100A9 in monocyte subsets specifically amplified in COVID-19 patients irrespectively of their ARDS status. These data suggest that, in the early stage of the disease, monocytes could contribute to the burst of circulating calprotectin (S100A8/S100A9), recently proposed to contribute to the secondary cytokine release syndrome described in severe COVID-19 and attributed to neutrophils (*18*).

Within severe COVID-19 patients, we detected no significant differences between ARDS^pos^ and ARDS^neg^ immune profiles, indicating a specificity of the phenotype induced by SARS-CoV2 infection, irrespectively of the respiratory complications. While most published studies showed differences between mild and severe COVID-19 diseases, some of their conclusions might be obscured by the fact that ARDS by itself, mechanical ventilation, and/or nonspecific treatments might impact immune parameters (*42*). A strength of our study comparing two groups of severe COVID-19 patients with or without ARDS is to highlight features directly related to the viral infection rather than to its respiratory complications or their treatment. Importantly, our cohort was homogeneous regarding treatment with in particular no immunosuppressive therapy at the time of sampling.

The small size of our cohort did not allow us to pinpoint a mortality prognostic factor based on our phenotypic data. However, we identified a specific immune pattern associated with the occurrence of the major adverse clinical events (thrombosis, nosocomial infection, septic shock, acute renal failure, and death) described in COVID-19 and combined as a clinical score. In particular, an increase of non-classical CD14^low^CD16^pos^ monocytes (Mo11), and CD14^pos^HLA-DR^low^ M-MDSC-like (Mo181), both not expressing CD169, are markers of adverse events. This suggests that besides the early increase of CD169^pos^ monocytes in all COVID-19 patients associated with T-cell dysfunctions, the immunological response to SARS-CoV2 infection features multiple alterations of monocytic subsets reflecting the severity of the disease. Consistent with these data, it was shown that CD14^pos^HLA-DR^low^ cells were increased in critical COVID-19 patients (*16*, *21*), while CD14^low^CD16^pos^ monocytes were correlated with the length of stay in ICU (*11*, *18*). To our knowledge, our study is the first to correlate the accumulation of non-classical monocytes and M-MDSCs occurring during the first days of ICU to adverse events.

Besides the low number of included patients, our study has some limitations. By focusing on severe patients with and without ARDS, we cannot make conclusions about phenotypic changes in mild and moderate diseases. Moreover, since the mass cytometry was conducted on PBMCs, we lack information on the neutrophil lineage, which appears affected in the COVID-19 (*18*). It would also be interesting to link these data with *in situ* data from lung tissue samples and bronchoalveolar lavages. However, our detailed analysis of circulating immune cells shows that immune monitoring of severe COVID-19 patients brings interesting prognostic biomarkers independently of their clinical classification in ARDS^pos^ versus ARDS^neg^. Moreover, we demonstrated that at the biological level, COVID-19 associated ARDS is different from other causes of ARDS, and might benefit from personalized therapy in addition to standard ARDS management (*18*, *43*).

## Materials and Methods

### Patients

This study was performed in the infectious diseases department and intensive care unit (ICU) at Rennes University Hospital. The study design was approved by our ethic committee (CHU Rennes, n°35RC20_9795_HARMONICOV, ClinicalTrials.gov Identifier: NCT04373200) and informed consent was obtained from patients in accordance with the Declaration of Helsinki. Peripheral blood was collected in tubes containing lithium heparin from COVID-19^neg^ARDS^pos^, COVID-19^pos^ARDS^pos^, and COVID-19^pos^ARDS^neg^ patients. Peripheral blood samples were drawn at D0 and D7. PBMC were isolated from whole blood using ficoll before cryopreservation. All patients provided written informed consent. The following data were recorded: gender, age, preexisting chronic kidney disease and acute kidney failure during the ICU stay (*44*), preexisting chronic heart failure (*45*), Body Mass Index (BMI), SAPS II at admission (*46*), duration of mechanical ventilation, length of hospital stay, and outcome (alive or dead) on day 7, day 30 and day 90. The occurrence of nosocomial infection, defined following CDC criteria as previously described (*47*), was also recorded during hospital stay. For each patient, a clinical score was built to summarize the occurrence of adverse clinical events frequently encountered during hospitalization (*47*, *48*). Each of the following events: thromboembolic events, nosocomial infection, septic shock, acute renal failure, and death counting as one point, the score varies from 0 (no adverse events) to 5. Patients characteristics are reported in table 1.

### Mass cytometry analysis

PBMC from patients were thawed. Briefly, cells were stained 5 minutes in RPMI supplemented with 0.5 μM Cisplatin Cell-ID™ (Fluidigm, San Francisco, CA) in RPMI 1640 before washing with 10% FCS in RPMI 1640. Cell pellets were resuspended in 80μl of 0.5% BSA in PBS. Then 60μl of each surface staining cocktail, lymphoid or myeloid, were added to 40μl of resuspended cells. After staining, cells were washed in 0.5% BSA in PBS before fixation/permeabilization with the transcription factor staining buffer set (Miltenyi, Bergisch-Gladbach, Germany). Then 60μl of each surface staining cocktail, lymphoid or myeloid, were added to 40μl of resuspended cells in Perm Buffer. After intracellular staining, cells were washed twice before staining in DNA intercalator solution (2.5% Paraformaldehyde, 1:3200 Cell-ID™ Intercalator-Ir (Fluidigm, San Francisco, CA) in PBS). Samples were cryopreserved at −80°C until acquisition on Helios™ System (Fluidigm, San Francisco, CA).

### CyTOF analysis pipeline

#### Pre-processing

After acquisition, intrafile signal drift was normalized and .fcs files were obtained using CyTOF software. To diminish batch effects, all files were normalized on EQ Beads (Fluidigm Sciences) using the premessa R package (https://github.com/ParkerICI/premessa). Files were then uploaded to the Cytobank cloud-based platform (Cytobank, Inc.). Data were first arcsinh-transformed using a cofactor of 5. For all files, live single cells were selected by applying a gate on DNA1 vs. DNA2 followed by a gate on DNA1 vs. Cisplatin, then beads were removed by applying a gate on the beads channel (Ce140Di) vs. DNA.1 Normalized, transformed and gated values were exported as FCS files.

#### CellCnn analysis

Identification of a Covid-19-specific cell-identity signature was carried out using the CellCnn algorithm (*28*), implemented in Pytorch in the ScaiVision platform (version 0.3.6, © Scailyte AG). Briefly, this is a supervised machine learning algorithm that trains a convolutional neural network with a single layer to predict sample-level labels using single-cell data as inputs.

The first 39 samples at day0 were analyzed by CellCnn. Data from each CyTOF panel was analyzed separately, in each case using all measured protein markers to train a series of CellCnn networks with varying hyperparameters. Each sample was given a label corresponding to the Covid-19 status of the patient from which the sample was drawn (positive or negative). To generate input data for training CellCnn, sub-samples of 2000 cells, termed multi-cell inputs (MCIs), were chosen randomly from each sample independently. For each training epoch, 2000 MCIs from each label class (Covid-19^pos^ or Covid-19^neg^) were presented to the network in random order. During training, 30 % of the samples were set aside for validation, chosen in a stratified manner to maintain the relative proportions of each class. 50 independent networks were generated for each CyTOF panel using hyperparameters randomly chosen from the following options: i) number of filters: (2, 3, 5, 7, and 10), ii) top-k pooling percentage: (1, 5, 10, 20, and 30), iii) dropout probability: (0.3, 0.4, and 0.6), iv) learning rate: (0.001, 0.003, and 0.01), and v) weight decay: (0.00001, 0.0001, 0.001, 0.01, and 0.1). Training was performed with a batch size of 50. Adam was used as an optimizer {kingma2015adam}, with a beta1 coefficient of 0.999 and a beta2 coefficient of 0.99. Each network was trained for a maximum of 50 epochs, or until the validation loss no longer decreased for 10 consecutive epochs. At the end of training, the weights from the epoch with lowest validation loss were returned. Representative filters were determined by clustering the filters from all networks achieving ≥ 90 % accuracy on the validation samples, then choosing the filter in each cluster with the minimum distance to all other filters in that cluster. For both CyTOF panels, a single representative filter showing the largest positive association with the Covid-19^pos^ label class was used to calculate cell-level filter response scores. Thresholds were set on the filter response scores to select Covid-19-associated cells by calculating the relative frequencies of selected cells in each sample at 100 different thresholds for each filter, then performing a logistic regression to predict sample labels. For each threshold, the data was first split in a stratified manner into a training set, comprising 60 % of samples, and a test set, comprising 40 % of samples. The logistic regression was performed on the training set, and the accuracy of resulting predictions was calculated on the test set. This procedure was performed 10 times, with randomly chosen training/test splits, and the mean of the resulting accuracies for each threshold was calculated. For the lymphoid panel, one threshold (9.63) achieved the highest accuracy and was set as the final threshold. For the myeloid panel, multiple thresholds achieved the same level of accuracy; the lowest of these (4.96) was set as the final threshold. The relative frequencies of cells in each sample with filter response scores greater than or equal to the respective thresholds were calculated and compared using a Wilcoxon rank-sum test.

#### viSNE, FlowSOM, and hierarchical clustering

We first performed a dimension reduction for both panels (i.e. myeloid and lymphoid) and all cleaned-up 63 files were first analyzed using viSNE, based upon the Barnes–Hut implementation of t-SNE. Equal downsampling was performed, based on the lowest event count in all files (lymphoid panel) or on the maximum total events allowed by Cytobank (myeloid panel). For the myeloid panel, the following parameters were used: perplexity = 45; iterations = 5000; theta = 0.5; all 37 channels selected. For the lymphoid panel the parameters were as follows: perplexity = 45; iterations = 7500; theta = 0.5; all 36 channels selected.

Then we applied a clustering method using the FlowSOM clustering algorithm. FlowSOM uses Self-Organizing Maps (SOMs) to partition cells into clusters based on their phenotype, and then builds a Minimal Spanning Tree (MST) to connect the nodes of the SOM, allowing the identification of metaclusters (i.e. group of clusters). We performed the FlowSOM algorithm on the previous viSNE results, using all events and panel channels, and the following parameters: clustering method = hierarchical consensus, iterations = 10, number of clusters = 256, number of metaclusters = 30. For both panels, each metacluster (containing a given number of clusters) was manually annotated based on his marker expression phenotype, his projection on the viSNE and his localization in the FlowSOM MST.

We first analyzed the myeloid panel. Out of 30 metaclusters defined by the FlowSOM approach, we identified 13 metaclusters with monocyte markers, other metaclusters contained other cell types, low count of cells or remaining doublets or dead cells. We visually identified 2 (Mo18 and Mo26) out or the 13 metaclusters that were heterogeneous. These 2 metaclusters were manually splited into 2 new metaclusters (identified respectively as Mo180, Mo181 and Mo214, Mo243) (Fig. S1B). Thus, altogether we analyzed 15 metaclusters of myeloid cells. Regarding the lymphoid compartment, we noticed that FlowSOM defined metaclusters at the lineage level, thus we retain all the 136 clusters included in 10 metaclusters of interest (i.e. containing lymphoid lineage markers) (Fig. S1C). All metaclusters and clusters phenotypes including their abundances and mean marker intensity were then exported from Cytobank for further analyses. Cytometry data was explored with Kaluza Analysis Software (Beckman Coulter). Hierarchical clustering and heatmaps were generated with R v3.6.3, using Rstudio v1.2.5033 and the pheatmap package.

#### Statistical analysis

Statistical analyses were performed with Graphpad Prism 8.4.3. P values were defined by a Kruskal-Wallis test followed by a Dunn’s post-test for multiple group comparisons or by Wilcoxon matched-pairs signed rank tests as appropriate. Correlations were calculated using Spearman test. * P < 0.05, ** P < 0.01, *** < 0.001, and **** P < 0.0001. Hierarchical clustering of the patients was performed using euclidean distance and complete clustering. Correspondence analysis was performed using the package factoshiny using as variable the abundance in cell subsets for each patient.

## Supplementary Materials

Figure S1: CyTOF experimental design and data analysis pipeline

Figure S2: CellCnn analysis (related to fig. 1B)

Figure S3: Abundance of Mo clusters (related to fig. 2C)

Figure S4: Heatmap of marker expression for the clusters from the lymphoid panel (related to fig. 3)

Figure S5: Correlation between myeloid metaclusters and lymphoid clusters (related to fig.3A)

Figure S6: Abundance of clusters from the lymphoid panel (related to fig. 3C)

Table S1: Panel of antibodies

Table S2: Spearman correlation between myeloid and lymphoid clusters

## Acknowledgments

We thank all donors, families, and surrogates, as well as the medical personnel in charge of patient care. We thank Catherine Blanc and Aurelien Corneau, from the CyPS core facility at Sorbonne University, Paris for access to the Helios mass cytometer.

## Author contributions

Conceptualization, M.R., F.R., M.Le, J.M.T., M.Cog., and K.T.; Methodology, M.R., S.L.G, J.D., and K.T; Formal analysis, M.R., J.Fer., S.L., and S.C.; Investigation, S.L.G., J.D., M.G., N.B., C.V., M.La., I.B., and M.Cor.; Ressources, F.R., M.Le., J.Feu., V.K.T., and J.M.T.; Data curation, M.R., J.Fer. and F.R.; Writing - original draft preparation, M.R. and J.Fer.; Writing - review and editing, M.R., J.Fer., S.L.G., S.C., V.K.T., J.M.T., M.Cog., and K.T.; Visualization, M.R. and J.Fer.; Supervision, M.R. and K.T.; Project administration, M.R. and K.T.; Funding acquisition, F.R. and M.Cog.;

## Data and materials availability

All data is available in the main text or the supplementary materials.

## References and Notes

1. E. J. Williamson, A. J. Walker, K. Bhaskaran, S. Bacon, C. Bates, C. E. Morton, H. J. Curtis, A. Mehrkar, D. Evans, P. Inglesby, J. Cockburn, H. I. McDonald, B. MacKenna, L. Tomlinson, I. J. Douglas, C. T. Rentsch, R. Mathur, A. Y. S. Wong, R. Grieve, D. Harrison, H. Forbes, A. Schultze, R. Croker, J. Parry, F. Hester, S. Harper, R. Perera, S. J. W. Evans, L. Smeeth, B. Goldacre, Factors associated with COVID-19-related death using OpenSAFELY, Nature 584, 430–436 (2020).

2. W.-J. Guan, Z.-Y. Ni, Y. Hu, W.-H. Liang, C.-Q. Ou, J.-X. He, L. Liu, H. Shan, C.-L. Lei, D. S. C. Hui, B. Du, L.-J. Li, G. Zeng, K.-Y. Yuen, R.-C. Chen, C.-L. Tang, T. Wang, P.-Y. Chen, J. Xiang, S.-Y. Li, J.-L. Wang, Z.-J. Liang, Y.-X. Peng, L. Wei, Y. Liu, Y.-H. Hu, P. Peng, J.-M. Wang, J.-Y. Liu, Z. Chen, G. Li, Z.-J. Zheng, S.-Q. Qiu, J. Luo, C.-J. Ye, S.-Y. Zhu, N.-S. Zhong, China Medical Treatment Expert Group for Covid-19, Clinical Characteristics of Coronavirus Disease 2019 in China, N Engl J Med 382, 1708–1720 (2020).

3. C. Huang, Y. Wang, X. Li, L. Ren, J. Zhao, Y. Hu, L. Zhang, G. Fan, J. Xu, X. Gu, Z. Cheng, T. Yu, J. Xia, Y. Wei, W. Wu, X. Xie, W. Yin, H. Li, M. Liu, Y. Xiao, H. Gao, L. Guo, J. Xie, G. Wang, R. Jiang, Z. Gao, Q. Jin, J. Wang, B. Cao, Clinical features of patients infected with 2019 novel coronavirus in Wuhan, China, Lancet 395, 497–506 (2020).

4. J. Helms, C. Tacquard, F. Severac, I. Leonard-Lorant, M. Ohana, X. Delabranche, H. Merdji, R. Clere-Jehl, M. Schenck, F. Fagot Gandet, S. Fafi-Kremer, V. Castelain, F. Schneider, L. Grunebaum, E. Anglés-Cano, L. Sattler, P.-M. Mertes, F. Meziani, CRICS TRIGGERSEP Group (Clinical Research in Intensive Care and Sepsis Trial Group for Global Evaluation and Research in Sepsis), High risk of thrombosis in patients with severe SARS-CoV-2 infection: a multicenter prospective cohort study, Intensive Care Med 46, 1089–1098 (2020).

5. R. Jeannet, T. Daix, R. Formento, J. Feuillard, B. François, Severe COVID-19 is associated with deep and sustained multifaceted cellular immunosuppression, Intensive Care Med, 1–3 (2020).

6. G. Chen, D. Wu, W. Guo, Y. Cao, D. Huang, H. Wang, T. Wang, X. Zhang, H. Chen, H. Yu, X. Zhang, M. Zhang, S. Wu, J. Song, T. Chen, M. Han, S. Li, X. Luo, J. Zhao, Q. Ning, Clinical and immunological features of severe and moderate coronavirus disease 2019, J Clin Invest 130, 2620–2629 (2020).

7. J. Villar, C. Ferrando, D. Martínez, A. Ambrós, T. Muñoz, J. A. Soler, G. Aguilar, F. Alba, E. González-Higueras, L. A. Conesa, C. Martín-Rodríguez, F. J. Díaz-Domínguez, P. Serna-Grande, R. Rivas, J. Ferreres, J. Belda, L. Capilla, A. Tallet, J. M. Añón, R. L. Fernández, J. M. González-Martín, dexamethasone in ARDS network, Dexamethasone treatment for the acute respiratory distress syndrome: a multicentre, randomised controlled trial, The Lancet Respiratory Medicine 8, 267–276 (2020).

8. Y.-N. Ni, G. Chen, J. Sun, B.-M. Liang, Z.-A. Liang, The effect of corticosteroids on mortality of patients with influenza pneumonia: a systematic review and meta-analysis, Crit Care 23, 99 (2019).

9. C. Agrati, A. Sacchi, V. Bordoni, E. Cimini, S. Notari, G. Grassi, R. Casetti, E. Tartaglia, E. Lalle, A. D’Abramo, C. Castilletti, L. Marchioni, Y. Shi, A. Mariano, J.-W. Song, J.-Y. Zhang, F.-S. Wang, C. Zhang, G. M. Fimia, M. R. Capobianchi, M. Piacentini, A. Antinori, E. Nicastri, M. Maeurer, A. Zumla, G. Ippolito, Expansion of myeloid-derived suppressor cells in patients with severe coronavirus disease (COVID-19), Cell Death Differ 395, 497 (2020).

10. P. S. Arunachalam, F. Wimmers, C. K. P. Mok, R. A. P. M. Perera, M. Scott, T. Hagan, N. Sigal, Y. Feng, L. Bristow, O. Tak-Yin Tsang, D. Wagh, J. Coller, K. L. Pellegrini, D. Kazmin, G. Alaaeddine, W.-S. Leung, J. M. C. Chan, T. S. H. Chik, C. Y. C. Choi, C. Huerta, M. Paine McCullough, H. Lv, E. Anderson, S. Edupuganti, A. A. Upadhyay, S. E. Bosinger, H. T. Maecker, P. Khatri, N. Rouphael, M. Peiris, B. Pulendran, Systems biological assessment of immunity to mild versus severe COVID-19 infection in humans, Science (2020), doi:10.1126/science.abc6261.

11. S. Chevrier, Y. Zurbuchen, C. Cervia, S. Adamo, M. E. Raeber, N. de Souza, S. Sivapatham, A. Jacobs, E. Bächli, A. Rudiger, M. Stüssi-Helbling, L. C. Huber, D. J. Schaer, J. Nilsson, O. Boyman, B. Bodenmiller, A distinct innate immune signature marks progression from mild to severe COVID-19, bioRxiv, 2020.08.04.236315 (2020).

12. C. Lucas, P. Wong, J. Klein, T. B. R. Castro, J. Silva, M. Sundaram, M. K. Ellingson, T. Mao, J. E. Oh, B. Israelow, T. Takahashi, M. Tokuyama, P. Lu, A. Venkataraman, A. Park, S. Mohanty, H. Wang, A. L. Wyllie, C. B. F. Vogels, R. Earnest, S. Lapidus, I. M. Ott, A. J. Moore, M. C. Muenker, J. B. Fournier, M. Campbell, C. D. Odio, A. Casanovas-Massana, Yale IMPACT Team, R. Herbst, A. C. Shaw, R. Medzhitov, W. L. Schulz, N. D. Grubaugh, C. Dela Cruz, S. Farhadian, A. I. Ko, S. B. Omer, A. Iwasaki, Longitudinal analyses reveal immunological misfiring in severe COVID-19, Nature 584, 463–469 (2020).

13. J. Hadjadj, N. Yatim, L. Barnabei, A. Corneau, J. Boussier, N. Smith, H. Pere, B. Charbit, V. Bondet, C. Chenevier-Gobeaux, P. Breillat, N. Carlier, R. Gauzit, C. Morbieu, F. Pene, N. Marin, N. Roche, T.-A. Szwebel, S. H. Merkling, J.-M. Treluyer, D. Veyer, L. Mouthon, C. Blanc, P.-L. Tharaux, F. Rozenberg, A. Fischer, D. Duffy, F. Rieux-Laucat, S. Kerneis, B. Terrier, Impaired type I interferon activity and inflammatory responses in severe COVID-19 patients, Science 31, eabc6027–15 (2020).

14. E. R. Mann, M. Menon, S. B. Knight, J. E. Konkel, C. Jagger, T. N. Shaw, S. Krishnan, M. Rattray, A. Ustianowski, N. D. Bakerly, P. Dark, G. Lord, A. Simpson, T. Felton, L. P. Ho, NIHR Respiratory TRC,, M. Feldmann, CIRCO,, J. R. Grainger, T. Hussell, Longitudinal immune profiling reveals key myeloid signatures associated with COVID-19, Sci. Immunol. 5, eabd6197 (2020).

15. D. Mathew, J. R. Giles, A. E. Baxter, D. A. Oldridge, A. R. Greenplate, J. E. Wu, C. Alanio, L. Kuri-Cervantes, M. B. Pampena, K. D’Andrea, S. Manne, Z. Chen, Y. J. Huang, J. P. Reilly, A. R. Weisman, C. A. G. Ittner, O. Kuthuru, J. Dougherty, K. Nzingha, N. Han, J. Kim, A. Pattekar, E. C. Goodwin, E. M. Anderson, M. E. Weirick, S. Gouma, C. P. Arevalo, M. J. Bolton, F. Chen, S. F. Lacey, H. Ramage, S. Cherry, S. E. Hensley, S. A. Apostolidis, A. C. Huang, L. A. Vella, UPenn COVID Processing Unit, M. R. Betts, N. J. Meyer, E. J. Wherry, Deep immune profiling of COVID-19 patients reveals distinct immunotypes with therapeutic implications, Science (2020), doi:10.1126/science.abc8511.

16. J. Schulte-Schrepping, N. Reusch, D. Paclik, K. Baßler, S. Schlickeiser, B. Zhang, B. Krämer, T. Krammer, S. Brumhard, L. Bonaguro, E. De Domenico, D. Wendisch, M. Grasshoff, T. S. Kapellos, M. Beckstette, T. Pecht, A. Saglam, O. Dietrich, H. E. Mei, A. R. Schulz, C. Conrad, D. Kunkel, E. Vafadarnejad, C.-J. Xu, A. Horne, M. Herbert, A. Drews, C. Thibeault, M. Pfeiffer, S. Hippenstiel, A. Hocke, H. Müller-Redetzky, K.-M. Heim, F. Machleidt, A. Uhrig, L. Bosquillon de Jarcy, L. Jürgens, M. Stegemann, C. R. Glösenkamp, H.-D. Volk, C. Goffinet, M. Landthaler, E. Wyler, P. Georg, M. Schneider, C. Dang Heine, N. Neuwinger, K. Kappert, R. Tauber, V. Corman, J. Raabe, K. M. Kaiser, M. T. Vinh, G. Rieke, C. Meisel, T. Ulas, M. Becker, R. Geffers, M. Witzenrath, C. Drosten, N. Suttorp, C. von Kalle, F. Kurth, K. Händler, J. L. Schultze, A. C. Aschenbrenner, Y. Li, J. Nattermann, B. Sawitzki, A.-E. Saliba, L. E. Sander, Deutsche COVID-19 OMICS Initiative (DeCOI), Severe COVID-19 Is Marked by a Dysregulated Myeloid Cell Compartment, Cell (2020), doi:10.1016/j.cell.2020.08.001.

17. T. Sekine, A. Perez-Potti, O. Rivera-Ballesteros, K. Strålin, J.-B. Gorin, A. Olsson, S. Llewellyn-Lacey, H. Kamal, G. Bogdanovic, S. Muschiol, D. J. Wullimann, T. Kammann, J. Emgård, T. Parrot, E. Folkesson, O. Rooyackers, L. I. Eriksson, J.-I. Henter, A. Sönnerborg, T. Allander, J. Albert, M. Nielsen, J. Klingström, S. Gredmark-Russ, N. K. Björkström, J. K. Sandberg, D. A. Price, H.-G. Ljunggren, S. Aleman, M. Buggert, Robust T cell immunity in convalescent individuals with asymptomatic or mild COVID-19, Cell, 1–46 (2020).

18. A. Silvin, N. Chapuis, G. Dunsmore, A.-G. Goubet, A. Dubuisson, L. Derosa, C. Almire, C. Hénon, O. Kosmider, N. Droin, P. Rameau, C. Catelain, A. Alfaro, C. Dussiau, C. Friedrich, E. Sourdeau, N. Marin, T.-A. Szwebel, D. Cantin, L. Mouthon, D. Borderie, M. Deloger, D. Bredel, S. Mouraud, D. Drubay, M. Andrieu, A.-S. Lhonneur, V. Saada, A. Stoclin, C. Willekens, F. Pommeret, F. Griscelli, L. G. Ng, Z. Zhang, P. Bost, I. Amit, F. Barlesi, A. Marabelle, F. Pene, B. Gachot, F. André, L. Zitvogel, F. Ginhoux, M. Fontenay, E. Solary, Elevated Calprotectin and Abnormal Myeloid Cell Subsets Discriminate Severe from Mild COVID-19, Cell (2020), doi:10.1016/j.cell.2020.08.002.

19. J.-W. Song, C. Zhang, X. Fan, F.-P. Meng, Z. Xu, P. Xia, W.-J. Cao, T. Yang, X.-P. Dai, S.-Y. Wang, R.-N. Xu, T.-J. Jiang, W.-G. Li, D.-W. Zhang, P. Zhao, M. Shi, C. Agrati, G. Ippolito, M. Maeurer, A. Zumla, F.-S. Wang, J.-Y. Zhang, Immunological and inflammatory profiles in mild and severe cases of COVID-19, Nat Commun 11, 3410 (2020).

20. A. J. Wilk, A. Rustagi, N. Q. Zhao, J. Roque, G. J. Martinez-Colon, J. L. McKechnie, G. T. Ivison, T. Ranganath, R. Vergara, T. Hollis, L. J. Simpson, P. Grant, A. Subramanian, A. J. Rogers, C. A. Blish, A single-cell atlas of the peripheral immune response in patients with severe COVID-19, Nat Med 26, 1070–1076 (2020).

21. E. J. Giamarellos-Bourboulis, M. G. Netea, N. Rovina, K. Akinosoglou, A. Antoniadou, N. Antonakos, G. Damoraki, T. Gkavogianni, M.-E. Adami, P. Katsaounou, M. Ntaganou, M. Kyriakopoulou, G. Dimopoulos, I. Koutsodimitropoulos, D. Velissaris, P. Koufargyris, A. Karageorgos, K. Katrini, V. Lekakis, M. Lupse, A. Kotsaki, G. Renieris, D. Theodoulou, V. Panou, E. Koukaki, N. Koulouris, C. Gogos, A. Koutsoukou, Complex Immune Dysregulation in COVID-19 Patients with Severe Respiratory Failure, Cell Host Microbe 27, 992–1000.e3 (2020).

22. M. Merad, J. C. Martin, Pathological inflammation in patients with COVID-19: a key role for monocytes and macrophages, Nat Rev Immunol 20, 355–362 (2020).

23. E. Z. Ong, Y. F. Z. Chan, W. Y. Leong, N. M. Y. Lee, S. Kalimuddin, S. M. Haja Mohideen, K. S. Chan, A. T. Tan, A. Bertoletti, E. E. Ooi, J. G. H. Low, A Dynamic Immune Response Shapes COVID-19 Progression, Cell Host Microbe 27, 879–882.e2 (2020).

24. S. De Biasi, D. Lo Tartaro, M. Meschiari, L. Gibellini, C. Bellinazzi, R. Borella, L. Fidanza, M. Mattioli, A. Paolini, L. Gozzi, D. Jaacoub, M. Faltoni, S. Volpi, J. Milić, M. Sita, M. Sarti, C. Pucillo, M. Girardis, G. Guaraldi, C. Mussini, A. Cossarizza, Expansion of plasmablasts and loss of memory B cells in peripheral blood from COVID-19 patients with pneumonia, Eur J Immunol 146, 3462 (2020).

25. S. De Biasi, M. Meschiari, L. Gibellini, C. Bellinazzi, R. Borella, L. Fidanza, L. Gozzi, A. Iannone, D. Lo Tartaro, M. Mattioli, A. Paolini, M. Menozzi, J. Milić, G. Franceschi, R. Fantini, R. Tonelli, M. Sita, M. Sarti, T. Trenti, L. Brugioni, L. Cicchetti, F. Facchinetti, A. Pietrangelo, E. Clini, M. Girardis, G. Guaraldi, C. Mussini, A. Cossarizza, Marked T cell activation, senescence, exhaustion and skewing towards TH17 in patients with COVID-19 pneumonia, Nat Commun 11, 3434 (2020).

26. N. Vabret, G. J. Britton, C. Gruber, S. Hegde, J. Kim, M. Kuksin, R. Levantovsky, L. Malle, A. Moreira, M. D. Park, L. Pia, E. Risson, M. Saffern, B. Salomé, M. E. Selvan, M. P. Spindler, J. Tan, V. van der Heide, J. K. Gregory, K. Alexandropoulos, N. Bhardwaj, B. D. Brown, B. Greenbaum, Z. H. Gümüş, D. Homann, A. Horowitz, A. O. Kamphorst, M. A. C. de Lafaille, S. Mehandru, M. Merad, R. M. Samstein, T. S. I. R. Project, M. Agrawal, M. Aleynick, M. Belabed, M. Brown, M. Casanova-Acebes, J. Catalan, M. Centa, A. Charap, A. Chan, S. T. Chen, J. Chung, C. C. Bozkus, E. Cody, F. Cossarini, E. Dalla, N. Fernandez, J. Grout, D. F. Ruan, P. Hamon, E. Humblin, D. Jha, J. Kodysh, A. Leader, M. Lin, K. Lindblad, D. Lozano-Ojalvo, G. Lubitz, A. Magen, Z. Mahmood, G. Martinez-Delgado, J. Mateus-Tique, E. Meritt, C. Moon, J. Noel, T. O’Donnell, M. Ota, T. Plitt, V. Pothula, J. Redes, I. R. Torres, M. Roberto, A. R. Sanchez-Paulete, J. Shang, A. S. Schanoski, M. Suprun, M. Tran, N. Vaninov, C. M. Wilk, J. Aguirre-Ghiso, D. Bogunovic, J. Cho, J. Faith, E. Grasset, P. Heeger, E. Kenigsberg, F. Krammer, U. Laserson, Immunology of COVID-19: Current State of the Science, Immunity 52, 910–941 (2020).

27. V. M. Ranieri, G. D. Rubenfeld, B. T. Thompson, N. D. Ferguson, E. Caldwell, E. Fan, L. Camporota, A. S. Slutsky, Acute respiratory distress syndrome: the Berlin Definition, JAMA 307, 2526–2533 (2012).

28. E. Arvaniti, M. Claassen, Sensitive detection of rare disease-associated cell subsets via representation learning, Nat Commun 8, 14825 (2017).

29. E. Galli, F. J. Hartmann, B. Schreiner, F. Ingelfinger, E. Arvaniti, M. Diebold, D. Mrdjen, F. Meer, C. Krieg, F. Al Nimer, N. Sanderson, C. Stadelmann, M. Khademi, F. Piehl, M. Claassen, T. Derfuss, T. Olsson, B. Becher, GM-CSF and CXCR4 define a T helper cell signature in multiple sclerosis, Nat Med, 1–27 (2019).

30. C. Krieg, M. Nowicka, S. Guglietta, S. Schindler, F. J. Hartmann, L. M. Weber, R. Dummer, M. D. Robinson, M. P. Levesque, B. Becher, High-dimensional single-cell analysis predicts response to anti-PD-1 immunotherapy, Nat Med 9, 2579–14 (2018).

31. S. Chevrier, J. H. Levine, V. R. T. Zanotelli, K. Silina, D. Schulz, M. Bacac, C. H. Ries, L. Ailles, M. A. S. Jewett, H. Moch, M. van den Broek, C. Beisel, M. B. Stadler, C. Gedye, B. Reis, D. Pe’er, B. Bodenmiller, An Immune Atlas of Clear Cell Renal Cell Carcinoma, Cell 169, 736–738.e18 (2017).

32. L. Kuri-Cervantes, M. B. Pampena, W. Meng, A. M. Rosenfeld, C. A. G. Ittner, A. R. Weisman, R. S. Agyekum, D. Mathew, A. E. Baxter, L. A. Vella, O. Kuthuru, S. A. Apostolidis, L. Bershaw, J. Dougherty, A. R. Greenplate, A. Pattekar, J. Kim, N. Han, S. Gouma, M. E. Weirick, C. P. Arevalo, M. J. Bolton, E. C. Goodwin, E. M. Anderson, S. E. Hensley, T. K. Jones, N. S. Mangalmurti, E. T. Luning Prak, E. J. Wherry, N. J. Meyer, M. R. Betts, Comprehensive mapping of immune perturbations associated with severe COVID-19, Sci. Immunol. 5, eabd7114 (2020).

33. M. J. Delano, P. A. Ward, The immune system’s role in sepsis progression, resolution, and longterm outcome, Immunol Rev 274, 330–353 (2016).

34. C. Ferrando, F. Suarez-Sipmann, R. Mellado-Artigas, M. Hernández, A. Gea, E. Arruti, C. Aldecoa, G. Martínez-Pallí, M. A. Martínez-González, A. S. Slutsky, J. Villar, COVID-19 Spanish ICU Network, Clinical features, ventilatory management, and outcome of ARDS caused by COVID-19 are similar to other causes of ARDS, Intensive Care Med 46, 846 (2020).

35. L. Gattinoni, S. Coppola, M. Cressoni, M. Busana, S. Rossi, D. Chiumello, COVID-19 Does Not Lead to a “Typical” Acute Respiratory Distress Syndrome, Am J Respir Crit Care Med 201, 1299–1300 (2020).

36. A.-S. Bedin, A. Makinson, M.-C. Picot, F. Mennechet, F. Malergue, A. Pisoni, E. Nyiramigisha, L. Montagnier, K. Bollore, S. Debiesse, D. Morquin, P. Bourgoin, N. Veyrenche, C. Renault, V. Foulongne, B. Caroline, B. Arnaud, V. Le Moing, P. Van de Perre, E. Tuaillon, Monocyte CD169 expression as a biomarker in the early diagnosis of COVID-19, medRxiv, 2020.06.28.20141556 (2020).

37. G. Carissimo, W. Xu, I. Kwok, M. Y. Abdad, Y.-H. Chan, S.-W. Fong, K. J. Puan, C. Y.-P. Lee, N. K.-W. Yeo, S. N. Amrun, R. S.-L. Chee, W. How, S. Chan, E. B. Fan, A. K. Andiappan, B. Lee, O. Rötzschke, B. E. Young, Y.-S. Leo, D. C. Lye, L. Renia, L. G. Ng, A. Larbi, L. F. P. Ng, Whole blood immunophenotyping uncovers immature neutrophil-to-VD2 T-cell ratio as an early prognostic marker for severe COVID-19, bioRxiv, 2020.06.11.147389 (2020).

38. A. Farina, G. Peruzzi, V. Lacconi, S. Lenna, S. Quarta, E. Rosato, A. R. Vestri, M. York, D. H. Dreyfus, A. Faggioni, S. Morrone, M. Trojanowska, G. A. Farina, Epstein-Barr virus lytic infection promotes activation of Toll-like receptor 8 innate immune response in systemic sclerosis monocytes, Arthritis Res Ther 19, 39 (2017).

39. H. Rempel, C. Calosing, B. Sun, L. Pulliam, F.-S. Wang, Ed. Sialoadhesin expressed on IFN-induced monocytes binds HIV-1 and enhances infectivity, PLoS ONE 3, e1967 (2008).

40. J. Carvelli, O. Demaria, F. Vély, L. Batista, N. C. Benmansour, J. Fares, S. Carpentier, M.-L. Thibult, A. Morel, R. Remark, P. André, A. Represa, C. Piperoglou, Explore COVID-19 IPH group, P. Y. Cordier, E. Le Dault, C. Guervilly, P. Simeone, M. Gainnier, Y. Morel, M. Ebbo, N. Schleinitz, E. Vivier, Explore COVID-19 Marseille Immunopole group, Association of COVID-19 inflammation with activation of the C5a-C5aR1 axis, Nature (2020), doi:10.1038/s41586-020-2600-6.

41. M. Liao, Y. Liu, J. Yuan, Y. Wen, G. Xu, J. Zhao, L. Cheng, J. Li, X. Wang, F. Wang, L. Liu, I. Amit, S. Zhang, Z. Zhang, Single-cell landscape of bronchoalveolar immune cells in patients with COVID-19, Nat Med 26, 842–844 (2020).

42. R. Stolk, E. van der Pasch, F. Naumann, J. Schouwstra, S. Bressers, T. van Herwaarden, J. Gerretsen, R. Schambergen, M. Ruth, H. van der Hoeven, H. van Leeuwen, P. Pickkers, M. Kox, Norepinephrine Dysregulates the Immune Response and Compromises Host Defense During Sepsis, Am J Respir Crit Care Med, rccm.202002–0339OC (2020).

43. E. Fan, J. R. Beitler, L. Brochard, C. S. Calfee, N. D. Ferguson, A. S. Slutsky, D. Brodie, COVID-19-associated acute respiratory distress syndrome: is a different approach to management warranted? The Lancet Respiratory Medicine 8, 816–821 (2020).

44. A. S. Levey, K.-U. Eckardt, Y. Tsukamoto, A. Levin, J. Coresh, J. Rossert, D. De Zeeuw, T. H. Hostetter, N. Lameire, G. Eknoyan, Definition and classification of chronic kidney disease: a position statement from Kidney Disease: Improving Global Outcomes (KDIGO), Kidney Int 67, 2089–2100 (2005).

45. P. Ponikowski, A. A. Voors, S. D. Anker, H. Bueno, J. G. F. Cleland, A. J. S. Coats, V. Falk, J. R. González-Juanatey, V.-P. Harjola, E. A. Jankowska, M. Jessup, C. Linde, P. Nihoyannopoulos, J. T. Parissis, B. Pieske, J. P. Riley, G. M. C. Rosano, L. M. Ruilope, F. Ruschitzka, F. H. Rutten, P. van der Meer, ESC Scientific Document Group, 2016 ESC Guidelines for the diagnosis and treatment of acute and chronic heart failure: The Task Force for the diagnosis and treatment of acute and chronic heart failure of the European Society of Cardiology (ESC)Developed with the special contribution of the Heart Failure Association (HFA) of the ESC. Eur Heart J 37, 2129–2200 (2016).

46. J. R. Le Gall, S. Lemeshow, F. Saulnier, A new Simplified Acute Physiology Score (SAPS II) based on a European/North American multicenter study, JAMA 270, 2957–2963 (1993).

47. B. Gaudriot, F. Uhel, M. Grégoire, A. Gacouin, S. Biedermann, A. Roisne, E. Flecher, Y. Le Tulzo, K. Tarte, J. M. Tadie, Immune Dysfunction After Cardiac Surgery with Cardiopulmonary Bypass: Beneficial Effects of Maintaining Mechanical Ventilation, Shock 44, 228–233 (2015).

48. P. Le Balc’h, K. Pinceaux, C. Pronier, P. Seguin, J. M. Tadie, F. Reizine, Herpes simplex virus and cytomegalovirus reactivations among severe COVID-19 patients, Crit Care 24, 530 (2020).

