## Supplemental Figures and Tables for "Mass cytometry and artificial intelligence define CD169 as a specific marker of SARS-CoV2-induced acute respiratory distress syndrome"

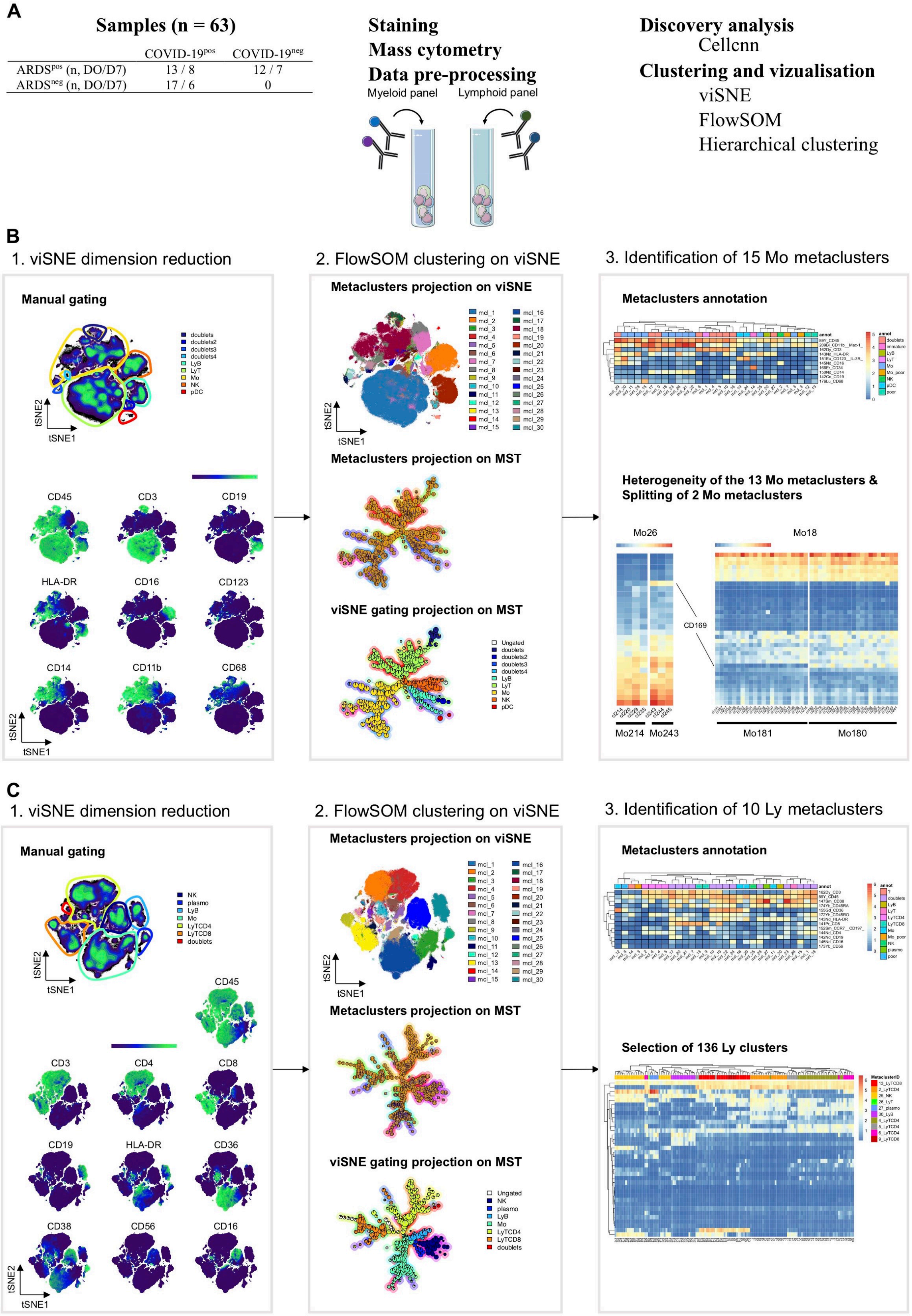
A
Samples (n = 63)
COVID-19pos
COVID-19neg
ARDSpos (n, DO/D7) 13 / 8 12 / 7
ARDSneg (n, DO/D7) 17 / 6 0
Staining
Mass cytometry
Data pre-processing
Myeloid panel
Lymphoid panel
Discovery analysis
Cellcnn
Clustering and vizualisation
viSNE
FlowSOM
Hierarchical clustering
B
1. viSNE dimension reduction
2. FlowSOM clustering on viSNE
3. Identification of 15 Mo metaclusters
Manual gating
doublets
doublets2
doublets3
doublets4
LyB
LyT
Mo
NK
pDC
tSNE2
tSNE1
CD45
CD3
CD19
HLA-DR
CD16
CD123
CD14
CD11b
CD68
Metaclusters projection on viSNE
Metaclusters projection on MST
viSNE gating projection on MST
Ungated
Metaclusters annotation
Heterogeneity of the 13 Mo metaclusters &
Splitting of 2 Mo metaclusters
Mo26
Mo18
CD169
Mo214
Mo243
Mo181
Mo180
C
1. viSNE dimension reduction
2. FlowSOM clustering on viSNE
3. Identification of 10 Ly metaclusters
Manual gating
NK
plasmo
LyB
Mo
LyTCD4
LyTCD8
doublets
CD45
CD3
CD4
CD8
CD19
HLA-DR
CD36
CD38
CD56
CD16
Metaclusters projection on viSNE
Metaclusters projection on MST
viSNE gating projection on MST
Metaclusters annotation
Selection of 136 Ly clusters

**Figure S1: CyTOF experimental design and data analysis pipeline**

**(A)** Experimental design for the processing of cryopreserved PBMC samples from COVID-19$^{neg}$ARDS$^{pos}$ (n = 12 at day 0 and n = 7 at day 7), COVID-19$^{pos}$ARDS$^{pos}$ (n = 13 at day O and n = 8 at day 7), and COVID-19$^{pos}$ARDS$^{neg}$ (n = 17 at day 0 and n = 6 at day 7) from staining to bioinformatics analysis. **(B)** Analysis pipeline for the myeloid panel allowing the identification of 15 monocytes metaclusters. Visualization of selected markers expression levels on a viSNE algorithm performed on all downsampled samples and manual gating (left). Metaclusters projection on viSNE and MST after clustering by FlowSOM, and visualization of manual gating on MST (middle). Manual annotation of 30 metaclusters determined by FlowSOM and identification of 15 monocytes metaclusters (Mo metaclusters). **(C)** Analysis pipeline for the lymphoid panel allowing the identification of relevant lymphocytes clusters. Visualization of selected markers expression levels on a viSNE algorithm performed on all downsampled samples and manual gating (left). Metaclusters projection on viSNE and MST after clustering by FlowSOM, and visualization of manual gating on MST (middle). Manual annotation of 30 metaclusters determined by FlowSOM and selection of the 136 lymphocytes corresponding clusters (Ly clusters) (right).

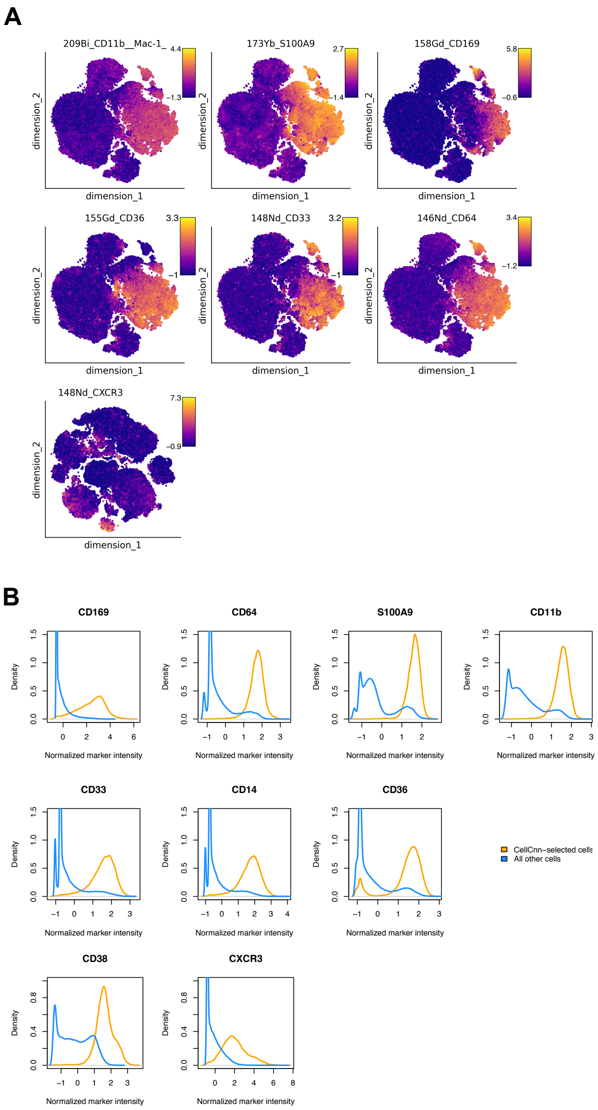


**Figure S2: CellCnn analysis (related to fig.1 B)**

CellCnn analysis performed on single cells from myeloid (top) and lymphoid panels (bottom) on 39 samples at Day 0 (COVID-19$^{neg}$ [n = 9] and COVID-19$^{pos}$ [n = 30]). **(A)** Cells defined by the best-performing CellCnn filters enrichment shown on tSNE and representative markers for each panel Representative markers for each panels. **(B)** Histogram of normalized marker intensity for the best-performing filter for each panel (orange) and the background (blue) for representative markers.

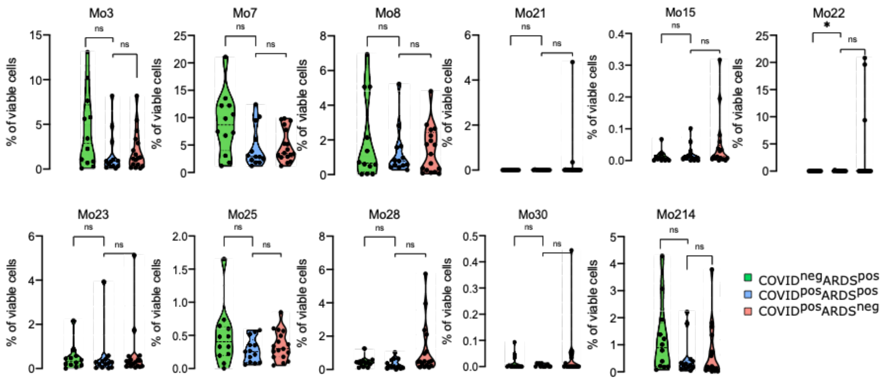


**Figure S3: Abundance of Mo clusters (related to fig. 2C)**

Abundance of metaclusters among singlet cells analyzed with the myeloid panel for COVID-19$^{neg}$ARDS$^{pos}$ (n=12, green), COVID-19$^{pos}$ARDS$^{pos}$ (n=13, blue), and COVID-19$^{pos}$ARDS$^{neg}$ (n=17, red). Kruskal-Wallis test with Dunn's multiple comparison correction.

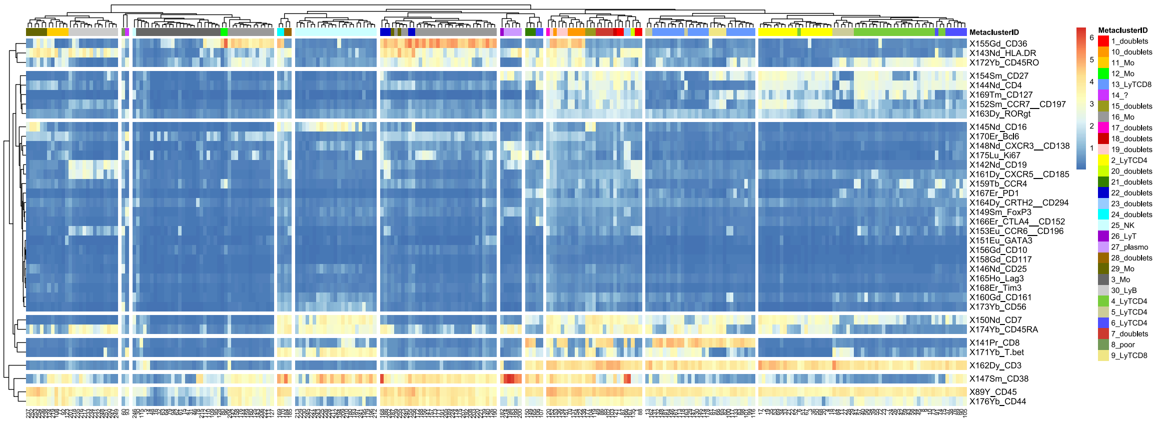


**Figure S4: Heatmap of marker expression for the clusters from the lymphoid panel (related to fig.3)**

Heatmap of all markers expression levels for all 256 clusters grouped into 30 metaclusters based on FlowSOM algorithm performed on the lymphoid panel, for COVID-19$^{neg}$ARDS$^{pos}$ (n=12, green), COVID-19$^{pos}$ARDS$^{pos}$ (n=13, blue), and COVID-19$^{pos}$ARDS$^{neg}$ (n=17, red).

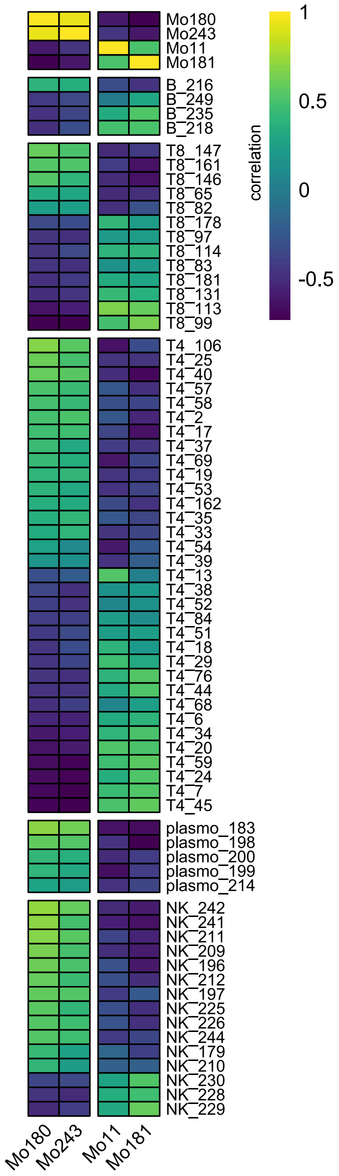


**Figure S5: Correlation between myeloid metaclusters and lymphoid clusters (related to fig.3A)**

Correlation between myeloid- (n = 4) and lymphoid- (n = 136) clusters from all patients at D0 (COVID-19$^{neg}$ARDS$^{pos}$ [n=12], COVID-19$^{pos}$ARDS$^{pos}$ [n=13], and COVID-19$^{pos}$ARDS$^{neg}$ [n=17]). Only significant Spearman correlation (P < 0.05) are shown (n = 70) (see table S2).

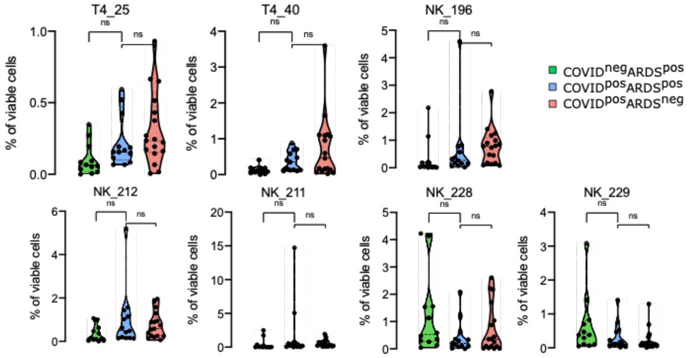


**Figure S6: Abundance of clusters from the lymphoid panel (related to fig. 3C)**

Comparison of abundance of lymphoid clusters between groups, among singlet cell analyzed. (COVID-19$^{neg}$ARDS$^{pos}$ [n=12], COVID-19$^{pos}$ARDS$^{pos}$ [n=13], and COVID-19$^{pos}$ARDS$^{neg}$ [n=17]). Kruskal-Wallis test with Dunn's multiple comparison correction.

### Table S1: Panel of antibodies

| | Myeloid | | Lymphoid | |
|---|---|---|---|---|
| Label | Target | Clone/Manufacturer | Target | Clone/Manufacturer |
| 89Y | CD45 | HI30/Fluidigm | CD45 | HI30/Fluidigm |
| 141Pr | CD326 | 9C4/BioLegend | CD8 | RPA-T8/BioLegend |
| 142Nd | CD19 | HIB19/BioLegend | CD19 | HIB19/BioLegend |
| 143Nd | HLA-DR | 10.1/BioLegend | HLA-DR | 10.1/BioLegend |
| 144Nd | CD31 | WM59/BioLegend | CD4 | RPA-T4/BioLegend |
| 145Nd | CD16 | B73.1/BioLegend | CD16 | B73.1/BioLegend |
| 146Nd | CD64 | L243/BioLegend | CD25 | BC96/BioLegend |
| 147Sm | CD11c | 3.9/BioLegend | CD38 | HIT2/BioLegend |
| 148Nd | CD33 | WM53/BioLegend | CXCR3 | G025H7/BioLegend |
| 149Sm | CD209 | 9E9A8/BioLegend | FoxP3 | 259D/C7 BD Biosciences |
| 150Nd | CD14 | M5E2/BioLegend | CD7 | CD7-6B7/BioLegend |
| 151Eu | CD123 | 6H6/BioLegend | GATA3 | TWAJ/Invitrogen |
| 152Sm | CD21 | Bu32/BioLegend | CCR7 | G043H7/BioLegend |
| 153Eu | CD192 | K036C2/BioLegend | CCR6 | G034E3/BioLegend |
| 154Sm | CD163 | GHI/61/BioLegend | CD27 | O323/BioLegend |
| 155Gd | CD36 | 5-271/BioLegend | CD36 | 5-271/BioLegend |
| 156Gd | CD86 | IT2.2/BioLegend | CD10 | HI10a/BioLegend |
| 158Gd | CD169 | 7.239/BioLegend | CD117 | 104D2/BioLegend |
| 159Tb | CD274 | 29E.2A3/BioLegend | CCR4 | L291H4/BioLegend |
| 160Gd | CD254 | MIH24/BioLegend | CD161 | HP-3G10/Biolegend |
| 161Dy | CD106 | EPR5047/Abcam | CXCR5 | J252D4/BioLegend |
| 162Dy | CD3 | UCHT1/BioLegend | CD3 | UCHT1/BioLegend |
| 163Dy | CD49a | TS2/7/BioLegend | RORgt | AFKJS-9/eBioscience |
| 164Dy | gp38 | REA446/Miltenyi Biotec | CRTH2 | BM16/BioLegend |
| 165Ho | CD80 | 2D10/BioLegend | LAG-3 | 7H2C65/BioLegend |
| 166Er | CD34 | 581/BioLegend | CTLA-4 | L3D10/BioLegend |
| 167Er | CD1a | HI149/BioLegend | PD1 | EH12.2H7/BioLegend |
| 168Er | CX3CR1 | 2A9-1/BioLegend | Tim-3 | F38-2E2/BioLegend |
| 169Tm | CD32 | FUN-2/BioLegend | CD127 | A019D5/BioLegend |
| 170Er | CD54 | HA58/BioLegend | Bcl-6 | k112-91/ BD Biosciences |
| 171Yb | CD195 | J418F1/BioLegend | T-bet | 4B10/BioLegend |
| 172Yb | CD206 | 15-2/BioLegend | CD45RO | UCHL1/BioLegend |
| 173Yb | S100A9 | A15105J/BioLegend | CD56 | HCD56/BioLegend |
| 174Yb | CD45RA | HI100/BioLegend | CD45RA | HI100/BioLegend |
| 175Lu | CD172a | 15-414/BioLegend | Ki67 | Ki-67/BioLegend |
| 176Yb | CD68 | Y1/82A/BioLegend | CD44 | BJ18/BioLegend |
| 209Bi | CD11b | ICRF44/Fluidigm | | |

**Table S2: Spearman correlation between myeloid and lymphoid clusters**

| row | column | cor | p | signif |
|---|---|---|---|---|
| Mo243 | Mo180 | 0.932 | 3.78E-11 | *** |
| NK_242 | Mo180 | 0.719 | 7.50E-05 | *** |
| plasmo_183 | Mo180 | 0.702 | 1.33E-04 | *** |
| NK_241 | Mo180 | 0.691 | 1.83E-04 | *** |
| NK_211 | Mo180 | 0.686 | 2.15E-04 | *** |
| T4_106 | Mo180 | 0.677 | 2.81E-04 | *** |
| NK_209 | Mo180 | 0.643 | 6.99E-04 | *** |
| NK_196 | Mo180 | 0.612 | 1.48E-03 | ** |
| T4_25 | Mo180 | 0.611 | 1.53E-03 | ** |
| T4_40 | Mo180 | 0.601 | 1.90E-03 | ** |
| NK_212 | Mo180 | 0.593 | 2.24E-03 | ** |
| T8_147 | Mo180 | 0.585 | 2.65E-03 | ** |
| plasmo_198 | Mo180 | 0.583 | 2.80E-03 | ** |
| NK_197 | Mo180 | 0.571 | 3.57E-03 | ** |
| NK_225 | Mo180 | 0.563 | 4.18E-03 | ** |
| T8_161 | Mo180 | 0.549 | 5.46E-03 | ** |
| NK_226 | Mo180 | 0.547 | 5.65E-03 | ** |
| NK_244 | Mo180 | 0.542 | 6.20E-03 | ** |
| T8_146 | Mo180 | 0.540 | 6.44E-03 | ** |
| NK_229 | Mo180 | -0.516 | 9.79E-03 | ** |
| NK_228 | Mo243 | -0.518 | 9.53E-03 | ** |
| T4_6 | Mo180 | -0.537 | 6.79E-03 | ** |
| Mo3 | Mo180 | -0.537 | 6.77E-03 | ** |
| T8_113 | Mo243 | -0.559 | 4.56E-03 | ** |
| T4_34 | Mo243 | -0.562 | 4.26E-03 | ** |
| T4_20 | Mo243 | -0.583 | 2.77E-03 | ** |
| Mo11 | Mo180 | -0.597 | 2.09E-03 | ** |
| T4_59 | Mo180 | -0.672 | 3.18E-04 | *** |
| T4_24 | Mo180 | -0.674 | 3.06E-04 | *** |
| T4_7 | Mo180 | -0.683 | 2.32E-04 | *** |
| T4_45 | Mo180 | -0.705 | 1.19E-04 | *** |
| T8_99 | Mo243 | -0.725 | 6.14E-05 | *** |
